# Unhealthy Yet Avoidable – How Cognitive Bias Modification Alters Behavioral And Brain Responses To Food Cues In Obesity

**DOI:** 10.1101/475020

**Authors:** Mehl Nora, Morys Filip, Villringer Arno, Horstmann Annette

**Affiliations:** Department of Neurology, Max Planck Institute for Human Cognitive and Brain Sciences, 04103 Leipzig, Germany; MaxNetAging Research School, 18057 Rostock, Germany; Leipzig University Medical Centre, IFB Adiposity Diseases, 04103 Leipzig, Germany; Leipzig University Medical Centre, Collaborative Research Centre 1052-A5, 04103 Leipzig, Germany

## Abstract

**Objective:** Obesity is associated with automatically approaching problematic stimuli, such as unhealthy food. Cognitive bias modification (CBM) could beneficially impact on problematic approach behavior. However, it is unclear which mechanisms are targeted by CBM in obesity: Candidate mechanisms include (1) altering reward value of food stimuli or (2) strengthening inhibitory abilities.

**Methods:** 33 obese people completed either CBM or sham training during fMRI scanning. CBM consisted of an implicit training to approach healthy and avoid unhealthy foods.

**Results:** At baseline, approach tendencies towards food were present in all participants. Avoiding vs. approaching food was associated with higher activity in the right angular gyrus (rAG). CBM resulted in a diminished approach bias towards unhealthy food, decreased activation in the rAG, and increased activation in the anterior cingulate cortex. Relatedly, functional connectivity between the rAG and right superior frontal gyrus increased. Analysis of brain connectivity during rest revealed training-related connectivity changes of the inferior frontal gyrus and bilateral middle frontal gyri.

**Conclusion:** Taken together, CBM strengthens avoidance tendencies when faced with unhealthy foods and alters activity in brain regions underpinning behavioral inhibition.

## 1. Introduction

The way we process and react to food cues is linked to unhealthy eating and obesity. Preferential processing and facilitated approach reactions towards food cues have been demonstrated in obese compared to healthy-weight individuals^1-4^. This automatic and biased processing may contribute to overconsumption of food^5,6^, especially in current ‘obesogenic’ environment.

Dual-process models address these automatic behavioral tendencies. The reflective-impulsive model^7^ states that during automatic behavior, the fast impulsive system overrules the slower reflective system. The former is guided by previously formed associations – approach positive and avoid negative stimuli – while the latter relies on explicit knowledge^7,8^. The incentive sensitization theory (e.g.^9^) further states that through repeated exposure, a reinforcer (e.g. tasty food) acquires incentive salience qualities via the brain’s reward system. In consequence, associated stimuli become attention-grabbing and the motivation to approach is increased^10^.

These theories can explain a paradox in maladaptive behaviors, where individuals continue to behave in disadvantageous ways despite better knowledge and often against personal intentions. Heavy drinkers, for example, were found to crave alcohol without necessarily liking it^11,12^ and were repeatedly found to display an approach bias towards alcohol^8,13,14^.

Cognitive bias modification (CBM) aims at changing these automatic tendencies towards problematic stimuli, subsequently improving maladaptive behavior. Promising effects have been shown in alcohol-dependent subjects and smokers^8,15,16^, where a CBM intervention decreased approach bias towards problematic stimuli while reducing consumption thereof. Regarding eating behavior, research has produced mixed findings^17^. Approach bias for and consumption of chocolate could be decreased through CBM in normal-weight individuals^18^. In contrast, no CBM effects were found in normal-weight females across three studies for both unhealthy food and chocolate^19^. In a sample of lean and obese participants, CBM reduced approach bias towards unhealthy food only in obese individuals^4^. Discrepancies in these results, however, might be explained by differences in samples and stimuli.

Despite the evidence for CBM’s efficacy in the behavioral context, not much research has been conducted regarding underlying neural mechanisms. One study investigated neural correlates of CBM in hazardous drinkers, where half of the participants received CBM while the other half received sham-training. CBM was associated with reducing activity in the medial prefrontal cortex^20^. This structure, together with the nucleus accumbens and the posterior cingulate gyrus, is engaged in approach behavior towards problematic stimuli^21,22^.

We investigated neural correlates of CBM in obesity by applying an approach-avoidance training^8^ in obese individuals in the fMRI scanner, randomly assigning participants to a training or a sham-training condition. The training was hypothesized to induce two effects: decreased approach tendencies towards unhealthy, and increased approach tendencies towards healthy foods. We hypothesized that CBM training could work through two mechanisms: by a) changing rewarding values of food stimuli and activation in reward-related regions; b) increasing inhibitory abilities and changing activity of brain regions engaged in inhibitory processing and cognitive control. We further explored whether CBM training induces differences in task-independent resting-state functional connectivity.

## 2. Materials and methods

### 2.1. Participants

33 obese participants (18-35 years) took part in the experiment (mean_BMI_=36.49kg/m^2^, σ=6.29, mean_age_=29.5 years, σ=4.5; for sample characteristics see Table S1). Sample size was selected according to a similar study investigating CBM effects in alcohol-dependent patients^22^. Participants met the following inclusion criteria: no history of neurological/psychological diseases, no thyroid disease, normal or regulated to normal blood pressure, no drug/alcohol addiction, no smoking, no MRI-related contraindications. Volunteers received a monetary compensation. The study was conducted according to the Declaration of Helsinki and approved by the Ethics Committee at the University of Leipzig. All participants gave written informed consent prior to the study.

### 2.2. Behavioral assessment

BMI was measured before the experiment. Before and after the MRI part, participants rated their mood, hunger and tiredness on visual analogue scales (VAS, 0-10). Subjects further sorted forty credit card-sized food pictures concerning their perceived healthiness and liking (see section: Selection of stimuli for details) on a cardboard with a scale (0-10). This picture set was independent of the one used in the fMRI task. It included healthy and unhealthy food pictures of comparable healthiness and liking to the fMRI picture set.

### 2.3. fMRI task

We used a training version of the approach-avoidance task (AAT; described in^4^, for task details see Figure 1A), which measures and modifies^4^ automatic approach and avoidance tendencies towards unhealthy and healthy food pictures (30 healthy/30 unhealthy). Participants react with push and pull movements of a joystick to the format of presented food pictures (e.g. push-vertical/pull-horizontal). The AAT consisted of three main phases: a *pre-phase*, a training or a sham-training phase, and a *post*-phase. In the *pre-, post*- and sham-training phases food pictures were presented equally often in push and pull formats. In the training phase 90% of unhealthy pictures appeared in a push format, and 90% of healthy pictures appeared in a pull format (Figure 1B). The amount of approach and avoidance responses was equal in both groups – any behavioral effects would therefore be related to the pictures’ content. We randomly assigned participants to a training or a sham-training group. Participants were not informed and not aware that training would take place, and the experiment introduction was performed by a blinded experimenter.

**Figure 1.**
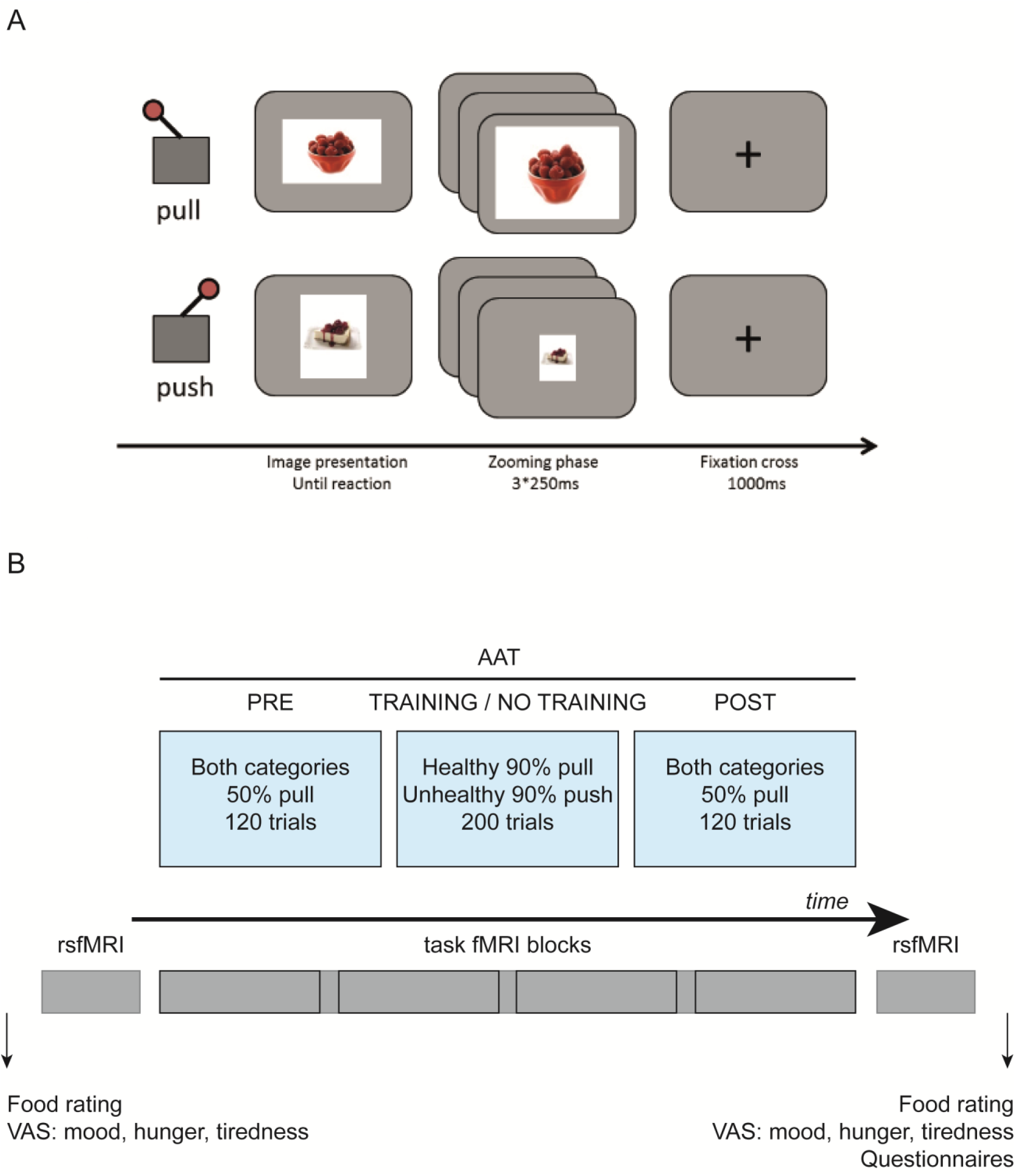
**A**: Modified Approach-Avoidance-Task. **B**: Overview of the experimental paradigm.

Further, we tested whether potential changes in automatic action tendencies are specific to pictures included in the training phase or generalized to the entire category (healthy vs. unhealthy). Therefore, only a subset of pictures used in *pre*- and *post-phases* was used for training (randomly chosen set of 20 out of 30 pictures for each participant).

The AAT lasted around 40 minutes and was symmetrically divided into four runs, each including 110 trials (independent of *pre*-, training, or *post*-phases). Participants were offered breaks between runs to relax and close their eyes.

### 2.4. Selection of stimuli

Food images for the AAT and the picture-sorting task were selected from the food-pics database^23^ and categorized into healthy and unhealthy according to^4^. Only images that were clearly identified as healthy or unhealthy were included^4^.

### 2.5. Neuroimaging

We collected resting-state (pre-AAT and post-AAT), task-related and anatomical neuroimaging data. Data were acquired using a 3T Siemens SKYRA scanner with a 20- channel head coil. For the AAT, 1104 T2*-weighted images were collected (TE=22ms, FA=90°, TR=2000ms, 40 slices, voxel size: 3.0×3.0×2.5mm^3^, distance factor: 20%, FoV: 192×192mm^2^, ascending order). 2*320 open-eyes resting-state T2*-weighted images were acquired using the same parameters. High-resolution anatomical MPRAGE image was acquired for each participant (TE=2.01ms, FA=9°, TR=2300ms, TI=900ms, voxel size: 1×1×1mm^3^, distance factor: 50%, FoV: 256×256mm^2^).

### 2.6. Data analyses

#### 2.6.1. Behavioral analysis

Mean reaction times were calculated for both picture categories (healthy/unhealthy food) and for both conditions (avoid/approach) during all three phases of the experiment. Bias scores were calculated as difference scores per category and condition: [healthy_push-healthy_pull] and [unhealthy_push-unhealthy_pull]. Positive scores reflect faster approach reactions for the respective food category, while negative scores indicate faster avoidance reactions.

No subject had to be excluded due to outliers or error rate. Outliers were defined as mean reaction times below or above 2 standard deviations from the group mean. The task was performed with high accuracy (mean = 97%, SD=3.23%).

During the *pre-phase,* bias scores significantly differing from zero would reflect baseline behavioral tendencies. Further, to ensure that no baseline group differences were present, we compared bias scores of training and the sham-training group. Analyses were carried out using one-sample and independent-samples t-tests, respectively.

Changes from *pre* to *post* were analyzed using a 2×2×2 rmANOVA. Group (training/sham-training) was used as a between-subject factor, and image category (healthy/unhealthy) and time (*pre/post*) as within-subject factors. We followed up by testing if bias scores significantly differed from zero in the *post-phase* with t-tests.

#### 2.6.2. fMRI data analysis

##### 2.6.2.1. Data preprocessing

Preprocessing of the MRI data included the following steps: skull-stripping of the anatomical data; motion correction, fieldmap correction, registration to anatomical data, slice-timing correction, and smoothing of the functional data. Further, white matter and cerebrospinal fluid signal were regressed out of the functional data. Finally, the functional data were registered to the MNI template. Prior to statistical analysis functional time series were high-pass filtered. For details see supplementary materials.

##### 2.6.2.2. AAT fMRI data analysis

A random-effects analysis was performed using SPM12. Regressors on a subject level were entered into a GLM and convolved with a double-gamma hemodynamic function. Contrast files were entered into second-level analysis, where we compared subjects as groups. BMI and age were entered into the analysis as covariates of no interest. By entering BMI as a covariate, we investigated general group effects dependent on training condition only and accounted for between-subject differences that could potentially be caused by differences in BMI. Age was entered as a covariate since groups differed in this respect. Results were thresholded at a whole-brain voxel-wise level with a threshold of 0.005 and on a cluster level with an FWE-corrected threshold of 0.05.

###### 2.6.2.2.1. GLM1: *Pre* and *post* data analysis

We investigated neural correlates of food approach and avoidance tendencies, as well as training effects on the brain. For the *pre-phase*, first level contrasts included general food approach and avoidance, and similar contrasts specific to each food category. We further compared food avoidance separately for healthy and unhealthy food pictures within and between groups. This was done to investigate whether CBM changes neural correlates underlying food approach and avoidance tendencies. To elucidate neural correlates of food approach tendencies independent of group membership, individual contrasts during the *pre-phase* of all participants were entered into a one-sample t-test on the second level. To ensure that no *pre*-training differences in task-related brain activity between groups were present, we entered individual contrasts into a two-sample t-test. Further, contrasts involving comparisons of *pre*- and *post-phases* were entered into two sample t-tests to investigate effects of training.

###### 2.6.2.2.2. GLM2: PPI analysis

To investigate whether CBM was related to changes in task-related functional connectivity, we conducted a psychophysiological interactions (PPI) analysis. It compares brain connectivity changes from a specified seed in the brain between two different experimental conditions. We placed the seed region in the right angular gyrus (6mm radius sphere) and tested whether connectivity of the seed in the unhealthy avoidance condition differed between experimental phases and groups.

##### 2.6.2.3. Resting-state fMRI data analysis

We acquired resting-state data to investigate whether effects of CBM are transferrable to functional changes outside the AAT. These data can be used to analyze resting-state functional connectivity – addressing how different brain regions interact. Resting-state connectivity analysis helps to understand how all brain regions generally interact with each other (degree centrality, DC), but also how specific *a priori* defined brain regions correlate with other brain areas (seed-based connectivity analysis, SCA).

To investigate connectivity in an exploratory fashion, but also in *a priori* defined regions, we used both the hypothesis-free approach of DC and the hypothesis-based approach of SCA with different inhibitory and reward-related seed regions – bilateral dorsolateral and medial prefrontal cortex (dlPFC and mPFC), middle frontal gyrus (MFG), bilateral amygdala, bilateral nucleus accumbens.

###### 2.6.2.3.1. Seed-based connectivity analysis

We investigated seed-based connectivity using predefined ROIs (see supplementary materials ‘regions of interest definition’) as seeds. Unsmoothed^24^ functional time series was entered into the analysis. Analysis was performed using Nipype and Nilearn algorithms. It resulted in sixteen connectivity maps - one for each ROI, and one for each phase of the experiment (*pre*- and *post*-AAT) - which were smoothed with a 6mm FWHM Gaussian kernel. *Pre*-AAT maps were subtracted from the *post*-AAT maps and resulting maps were entered into a two-sample t-test to investigate group differences using FWE-corrected and Bonferroni corrected (number of seeds) statistical thresholds.

###### 2.6.2.3.2. Degree centrality

Degree centrality denotes a number of direct connections of a node to all other nodes in the network^25^. We used AFNI within the Nipype framework, with correlation thresholds of 0.5 (only r>0.5 included in results). Firstly, we calculated DC maps separately for each participant and phase of the experiment. Secondly, similarly to SCA analysis, we subtracted the *pre*-AAT maps from the *post* AAT maps and entered resulting volumes into a two-sample t-test.

## 3. Results

### 3.1. Behavioral results

There were no baseline group differences regarding hunger, tiredness and mood (smallest p=.534, Table S1), and subjective ratings of the AAT stimulus material (smallest p=.498, Table S3). All participants rated healthy images as healthier (t(32)=36.723; p<.001, d=6.40) and liked them more (t(32)=5.507; p<.001. d=0.96) than unhealthy images.

To test if CBM would affect explicit ratings of the independent picture set of the sorting task (Table S4), we performed separate 2×2×2 rmANOVAs for healthiness and liking ratings with image category, time and group as factors. They revealed no interactions with group (healthiness: F(1,31)=0.001, p=0.975, η^2^p=.000; liking: F(1,31)=0.453, p=0.503, η^2^p=.014), indicating no effects of CBM on this task. This was confirmed by an Bayesian rmANOVA (JASP version 0.9; JASP Team 2018) with identical factors – lack of differential group effect for healthiness and liking ratings was 2.886 and 3.847 times more likely than the alternative explanation, respectively.

#### 3.1.1. AAT

We first tested for baseline food approach bias. As expected, both groups showed a significant food approach bias, as bias scores for healthy and unhealthy images differed from zero (training group: t(16)=2.994, p=.009 and t(16)=2.334, p=.033, sham-training group: t(15)=3.728, p=.002 and t(15)=2.218, p=.042, respectively). There were no group differences in approach bias (healthy: t(31)=0.557, p=.581, unhealthy: t(31)=-0.465, p=0.645).

Our main question was whether CBM affects approach behavior towards food stimuli and whether this depends on picture category. We used a 2×2×2 rmANOVA with group, image category and time as factors. We found a significant three-way interaction of group (training or sham-training), image category (healthy vs. unhealthy) and time (*pre*- vs. *post*- training; F(1,31)=8.902, p=.006, η^2^p=.223). Follow-up paired t-tests indicated that training group, as opposed to sham-training group, decreased approach tendencies towards unhealthy images (Table 1, Figure 2), which was driven by significantly faster push movements for unhealthy images (t(16)=2.735, p=.015).

**Table 1.**
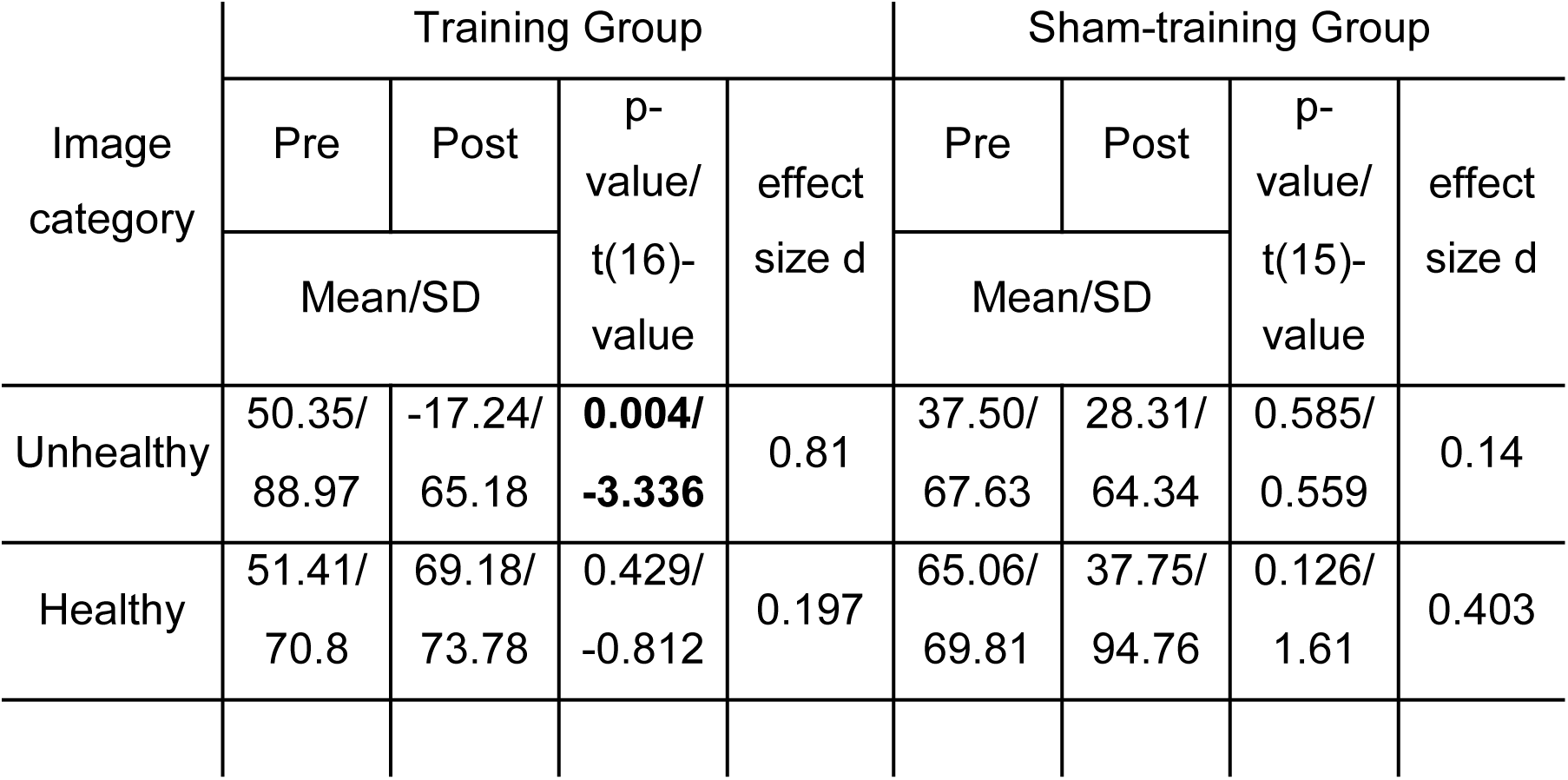
Bias scores for healthy and unhealthy images in the training and sham-training group for the *pre*- and *post*-phases. P-values reflect significance of changes from *pre*- to *post*- in bias scores.

**Figure 2.**
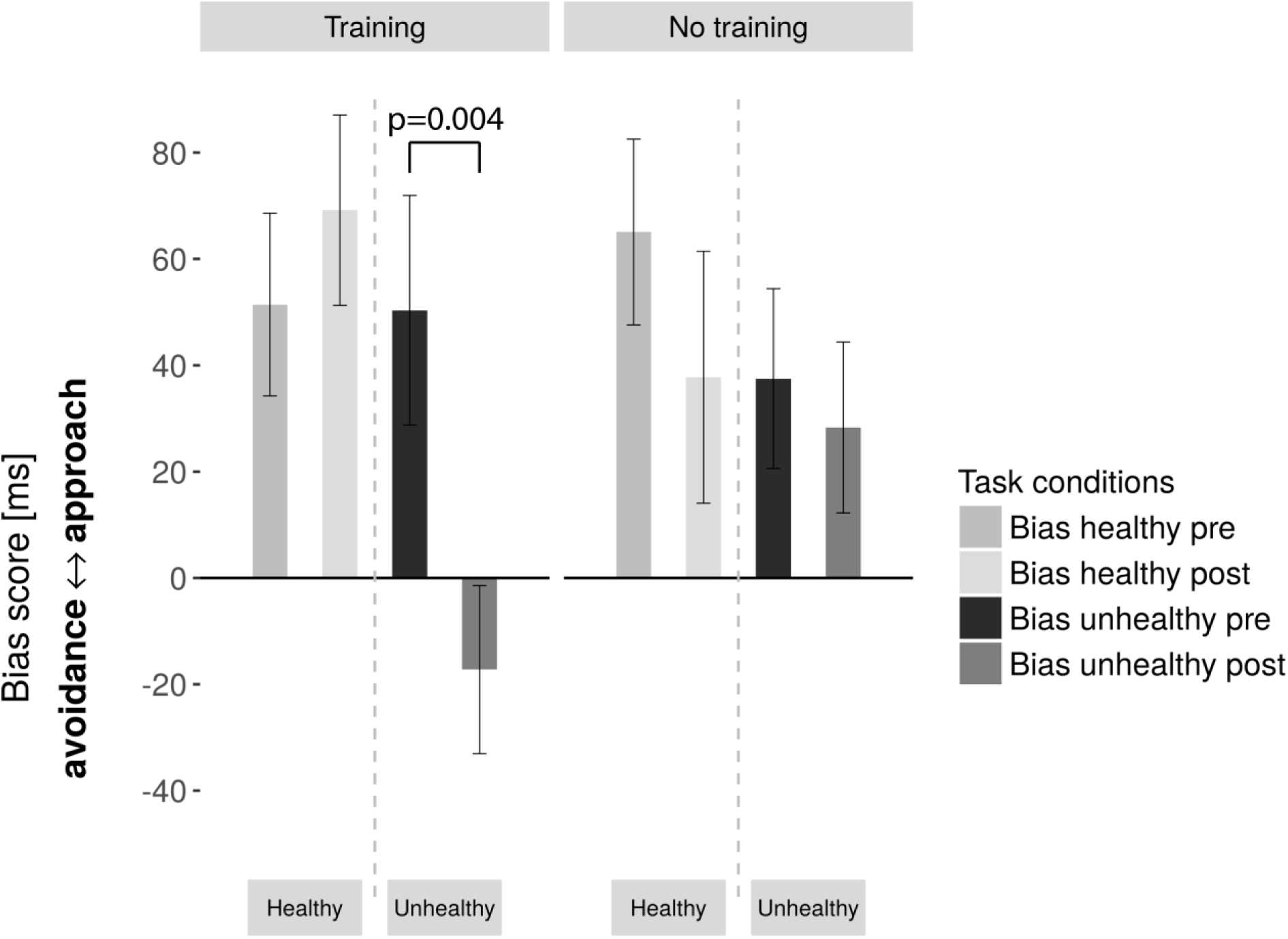
Bias scores in the training and sham-training groups *pre*- and *post*-training (error bars: standard error of the mean). We observed a significant three-way interaction between group, time, and image category.

To test for generalization, a 2×2×2 rmANOVA was performed in the training group. Factors included picture set (trained vs. not-trained images), image category and time. A significant three-way interaction would indicate lack of generalization, showing that bias scores for trained and not-trained images of the same category were not similarly affected by the training. The three-way interaction was marginally not significant (F(1,15)=4.464, p=.051, η^2^p=.218), suggesting that generalization might have occurred. A follow-up Bayesian rmANOVA with identical factors showed that a model with the three-way interaction, compared to a model without this interaction was 0.493 less likely, indicating anecdotal evidence in favor of generalization. We therefore cannot conclude with certainty, whether generalization occurred.

### 3.2. Neuroimaging results

#### 3.2.1. GLM1

##### 3.2.1.1. Baseline food approach and avoidance

We investigated neural correlates of *pre*-training approach and avoidance tendencies with a one-sample t-test, as in this stage groups did not differ in any way. Contrasts included food approach and food avoidance, together and separately for healthy and unhealthy food. A contrast corresponding to general food avoidance (push>pull independent of picture category) revealed significant clusters in the right angular gyrus (rAG) and the cuneus (Figure 3A). Food avoidance activations were driven by the unhealthy food category (push>pull for unhealthy food). For the general approach for food (pull>push), we found a significant cluster in the left postcentral gyrus (Figure 3B, Table 2). We found no further significant results and did not find group differences for above-mentioned contrasts.

**Table 2.**
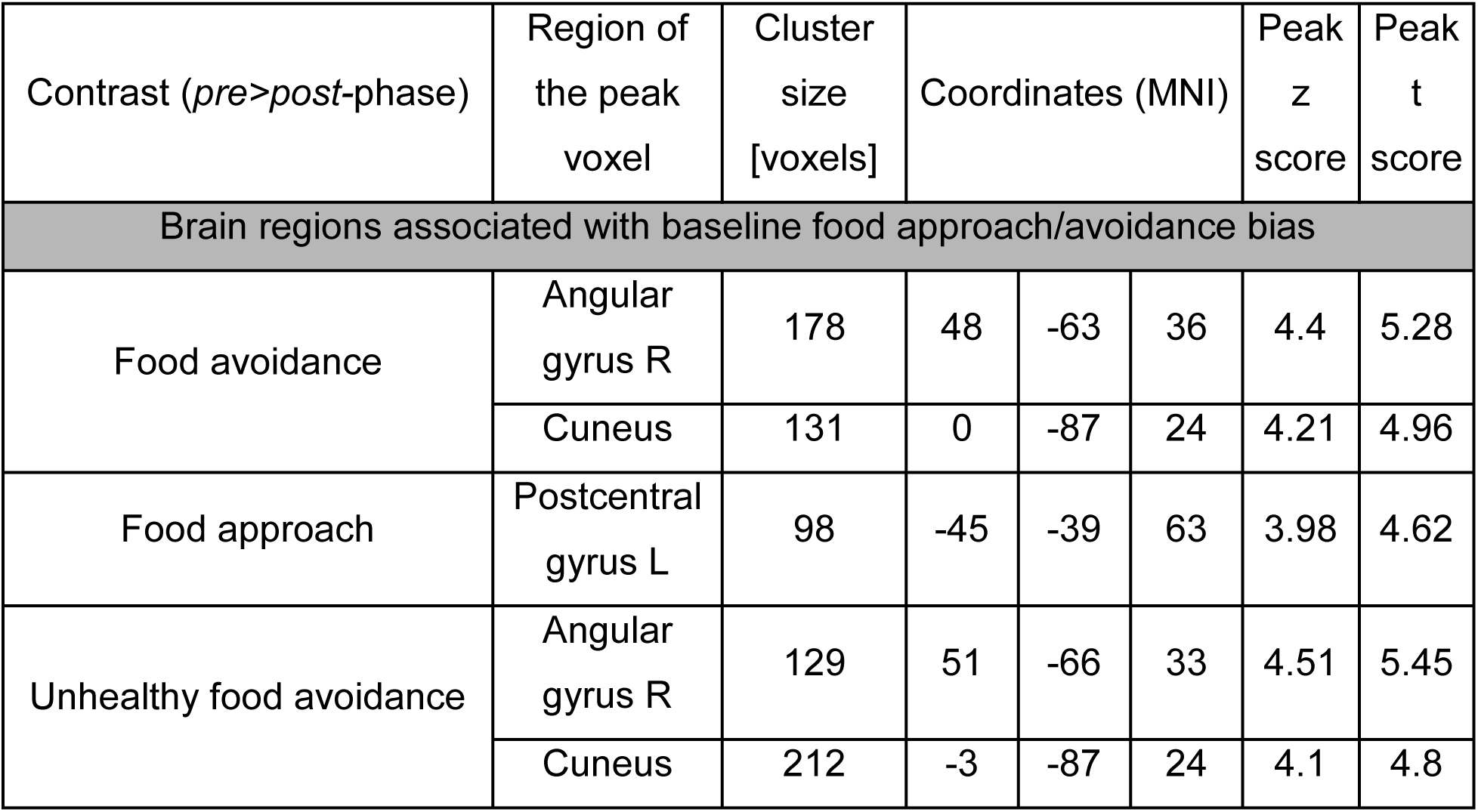
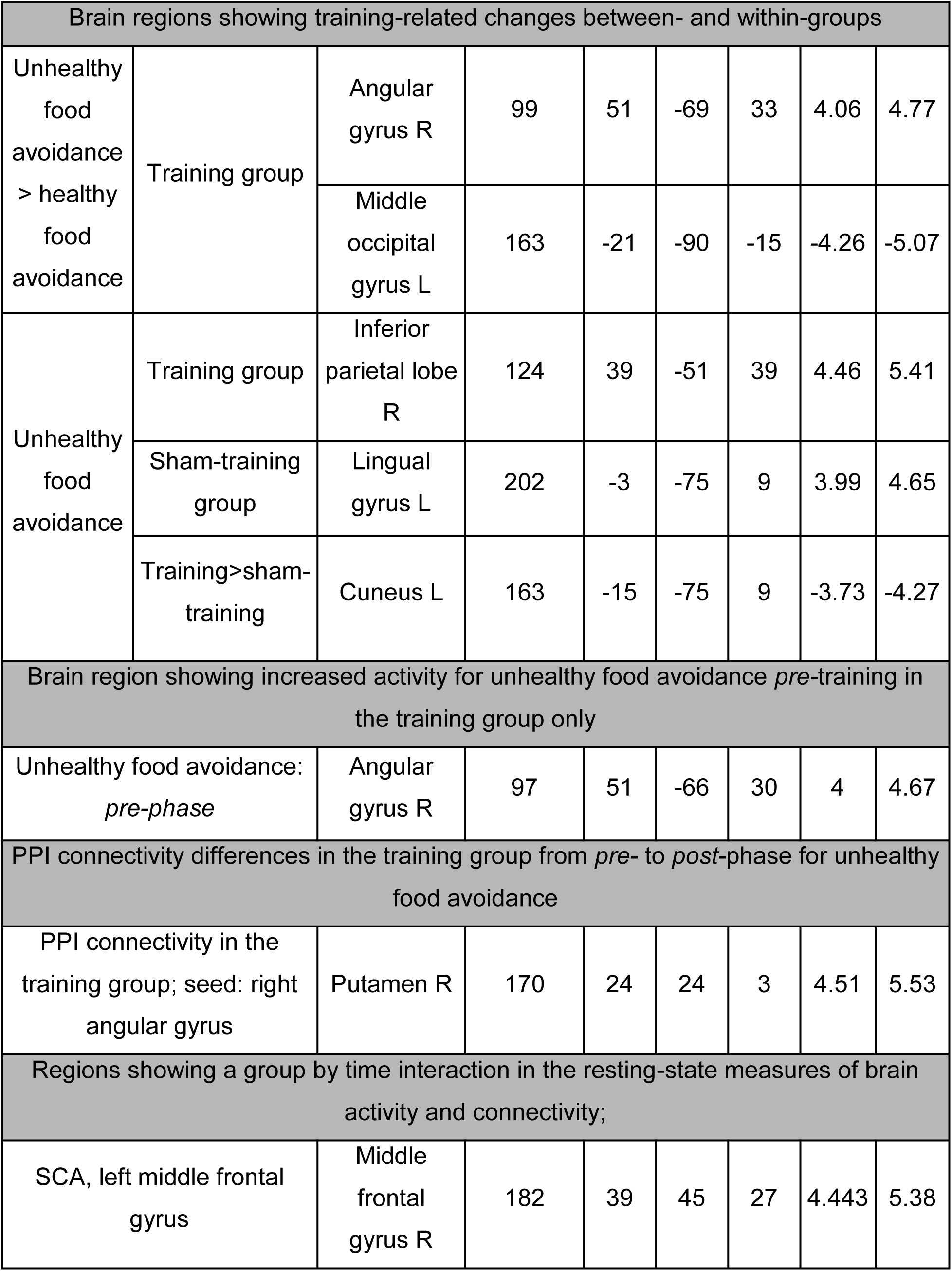
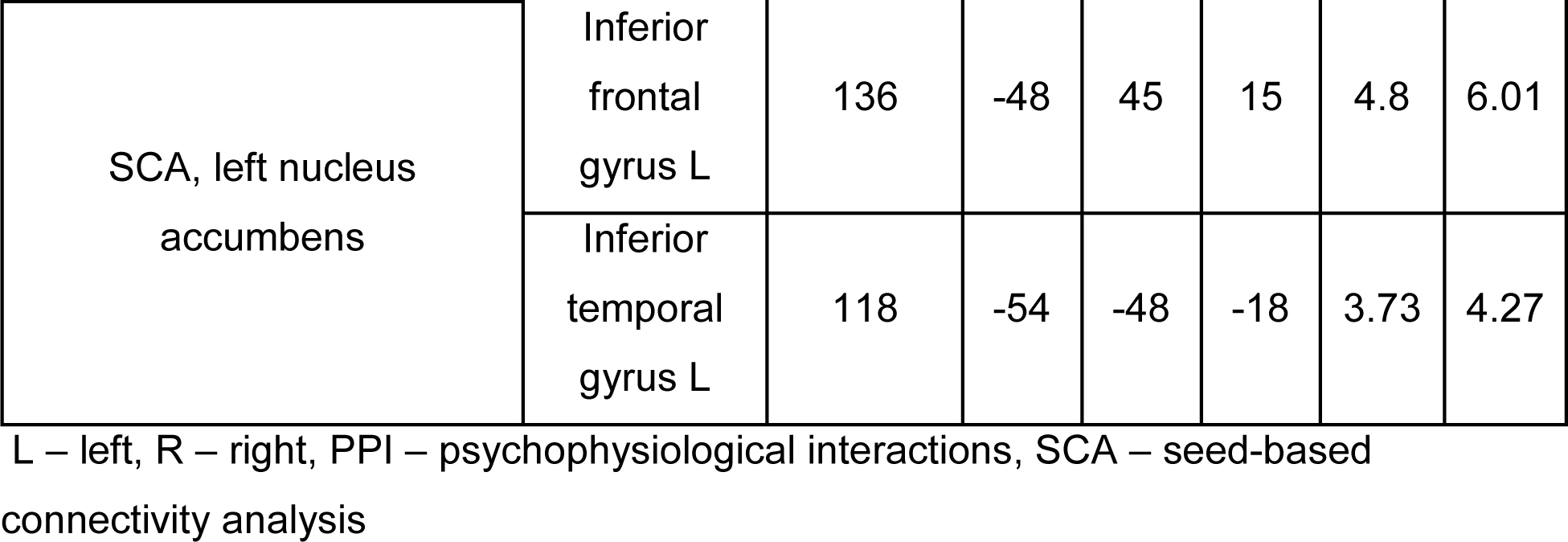
Brain regions showing training-related changes between- and within-groups

| Contrast (*pre>post*-phase) | Region of the peak voxel | Cluster size [voxels] | Coordinates (MNI) |  |  | Peak z score | Peak t score |
| --- | --- | --- | --- | --- | --- | --- | --- |
| Brain regions associated with baseline food approach/avoidance bias |  |  |  |  |  |  |  |
| Food avoidance | Angular gyrus R | 178 | 48 | -63 | 36 | 4.4 | 5.28 |
|  | Cuneus | 131 | 0 | -87 | 24 | 4.21 | 4.96 |
| Food approach | Postcentral gyrus L | 98 | -45 | -39 | 63 | 3.98 | 4.62 |
| Unhealthy food avoidance | Angular gyrus R | 129 | 51 | -66 | 33 | 4.51 | 5.45 |
|  | Cuneus | 212 | -3 | -87 | 24 | 4.1 | 4.8 |

|  |  |  |  |  |  |  |  |  |
| --- | --- | --- | --- | --- | --- | --- | --- | --- |
| Unhealthy food avoidance > healthy food avoidance | Training group | Angular gyrus R | 99 | 51 | -69 | 33 | 4.06 | 4.77 |
|  |  | Middle occipital gyrus L | 163 | -21 | -90 | -15 | -4.26 | -5.07 |
| Unhealthy food avoidance | Training group | Inferior parietal lobe R | 124 | 39 | -51 | 39 | 4.46 | 5.41 |
|  | Sham-training group | Lingual gyrus L | 202 | -3 | -75 | 9 | 3.99 | 4.65 |
|  | Training>sham-training | Cuneus L | 163 | -15 | -75 | 9 | -3.73 | -4.27 |
| Brain region showing increased activity for unhealthy food avoidance *pre*-training in the training group only |  |  |  |  |  |  |  |  |
| Unhealthy food avoidance: *pre-phase* |  | Angular gyrus R | 97 | 51 | -66 | 30 | 4 | 4.67 |
| PPI connectivity differences in the training group from *pre*- to *post*-phase for unhealthy food avoidance |  |  |  |  |  |  |  |  |
| PPI connectivity in the training group; seed: right angular gyrus |  | Putamen R | 170 | 24 | 24 | 3 | 4.51 | 5.53 |
| Regions showing a group by time interaction in the resting-state measures of brain activity and connectivity; |  |  |  |  |  |  |  |  |
| SCA, left middle frontal gyrus |  | Middle frontal gyrus R | 182 | 39 | 45 | 27 | 4.443 | 5.38 |
| SCA, left nucleus accumbens | Inferior frontal gyrus L | 136 | -48 | 45 | 15 | 4.8 | 6.01 |
|  | Inferior temporal gyrus L | 118 | -54 | -48 | -18 | 3.73 | 4.27 |

L – left, R – right, PPI – psychophysiological interactions, SCA – seed-based connectivity analysis

**Figure 3.**
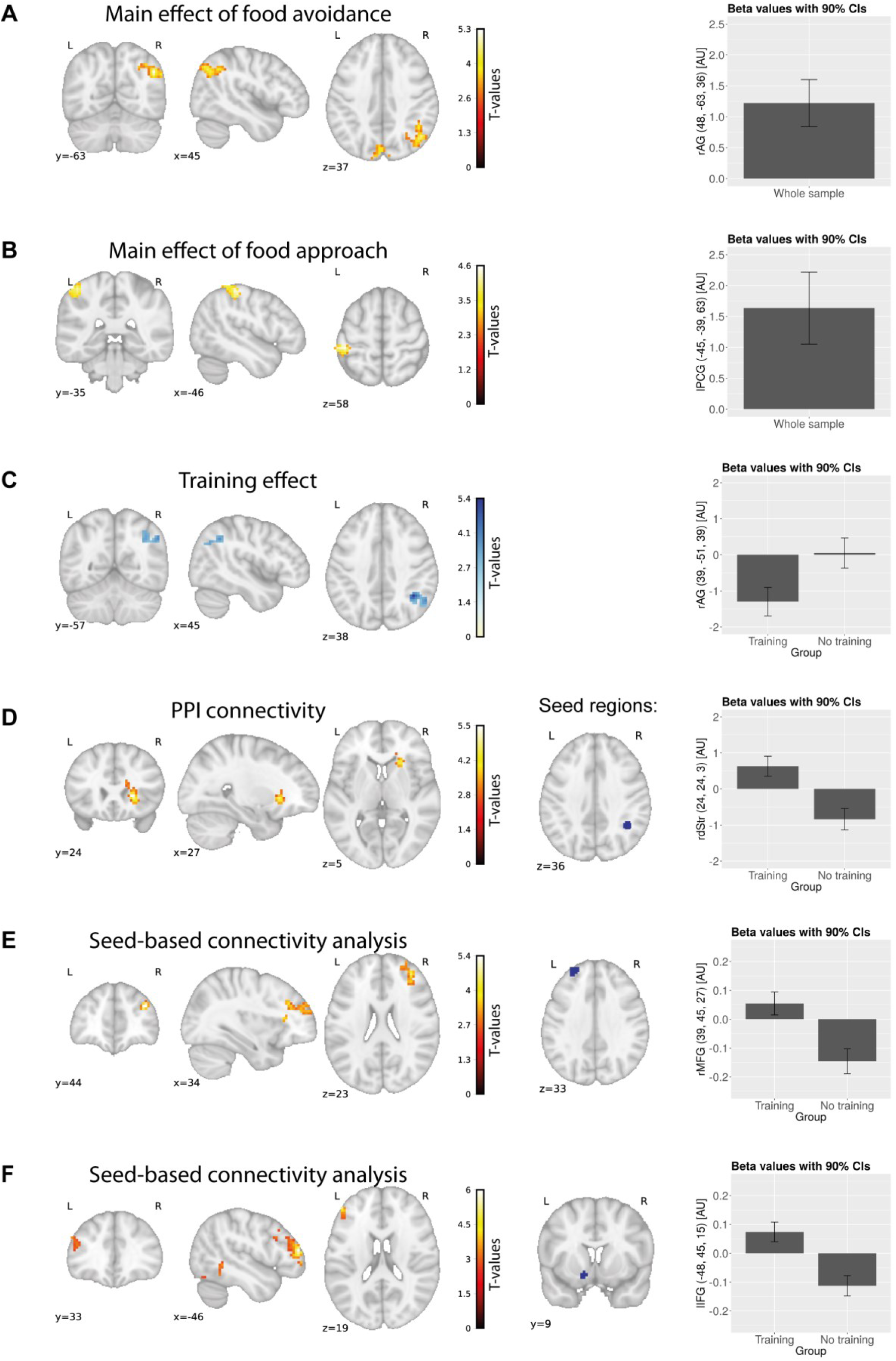
Main effects of food approach/avoidance *pre*-training in both groups together (A&B), and effects of CBM training along with contrast estimates (arbitrary units - AU); note a different scale in the parameter estimates for subfigures. **A:** Main effect of food avoidance. **B:** Main effect of food approach. **C:** Training effect was reflected in a decreased brain activity in the right angular gyrus for healthy food avoidance vs. unhealthy food avoidance. **D:** Higher task-related connectivity between the right dorsal striatum and the right angular gyrus in the unhealthy push vs. unhealthy pull condition after training. **E:** Increased connectivity *post*-training in the training vs. sham-training group in the left and right middle frontal gyri (resting-state seed-based connectivity analysis). **F:** Increased connectivity *post*-training in the training vs. sham-training group in the left nucleus accumbens and inferior frontal gyrus (resting-state seed-based connectivity analysis). rAG – right angular gyrus, lPCG - left postcentral gyrus, rdStr – right dorsal striatum, rMFG – right middle frontal gyrus, lIFG – left inferior frontal gyrus, AU – arbitrary units, CI – confidence intervals.

##### 3.2.1.2. *Pre*- to *post*- changes

As the main analysis of interest, we tested whether the training effect – decreased approach bias towards unhealthy foods – was associated with neuronal changes. We contrasted unhealthy food avoidance (unhealthy_push>unhealthy_pull) with healthy food avoidance (healthy_push>healthy_pull) before vs. after training. In the training group, the rAG showed decreased activity *post*-training, whereas the left middle occipital gyrus showed increased activity (Figure 3). We then compared unhealthy food conditions (unhealthy_push>unhealthy_pull) *pre*- and *post*-training and found a similar effect. Here, we also found decreased activity in the left lingual gyrus for the sham-training group, and a group difference in the cuneus (training>sham-training group). The effect in the rAG was driven by lower brain activity in the training group for the *post- vs. pre-phase* for unhealthy food avoidance (unhealthy_push>unhealthy_pull). Results indicate that brain activity in the right rAG for pushing unhealthy foods in the training group decreased after the training (Table 2).

##### 3.2.1.3. PPI analysis

We consistently found the rAG to be associated with unhealthy food avoidance and the effects of CBM, and therefore performed PPI analysis with the rAG as the seed. We compared connectivity differences for unhealthy food avoidance between *pre*- and *post-phases*. This analysis showed a significant cluster in the rSFG/rMFG and in the right caudate/putamen, indicating that connectivity between the rAG and these structures increased *post*-training in the training group (Table 2, Figure 3).

#### 3.2.2. Resting-state data

##### 3.2.2.1. Seed-based connectivity analysis

We investigated whether training induced connectivity changes between previously specified seed regions using resting-state, task-independent data. For the left MFG, we found a significant group*time interaction in the right MFG. We observed a similar interaction effect for connectivity between the left nucleus accumbens and the left inferior frontal gyrus (IFG; Table 2, Figure 3).

##### 3.2.2.2. Degree centrality

Similar to SCA, DC describes task-independent connectivity changes within the brain. These changes, however, are general and not specific to chosen ROIs. In our study this analysis did not produce any significant results.

## 4. Discussion

We investigated the underlying neural mechanisms of CBM in obese individuals. To this end, a training form of the AAT was applied in the fMRI scanner, where half of the participants received training, while the other half underwent sham training. This between-group design combined with fMRI measures allowed us to clarify whether the effects of CBM are mediated by a) changing rewarding values of food stimuli and brain activation in reward-related brain regions, or b) increasing inhibitory abilities and affecting brain regions engaged in inhibitory processing and cognitive control. We found that all participants showed faster approach than avoidance reactions towards healthy and unhealthy food images, suggesting that approaching food is an automatic process. This was paralleled by our findings on the neural level, where the rAG showed higher activation for avoiding food – a potentially conflicting situation. The rAG is a part of the temporoparietal junction (TPJ), which is often related to both processing of social cues and attentional processes^26,27^. CBM specifically affected the training group, where approach tendencies towards unhealthy food were successfully decreased. This was related to a lower activation in the rAG after training. Additionally, we observed group-specific changes in resting-state connectivity between inhibitory regions, such as the MFG or the IFG^28-31^, and in task-related connectivity between the rAG and the right caudate/putamen (dorsal striatum). Avoiding food thus appears to be a potentially conflicting situation, requiring activation of inhibitory and conflict resolution brain mechanisms. CBM seems to decrease this demand by means of strengthening connectivity between inhibitory brain regions. Further, we found no evidence for altered reward valuation of food stimuli after CBM in both behavioral and imaging data.

As mentioned, avoiding food was related to higher activity of the rAG – a part of the TPJ. Bzdok and colleagues showed that the right TPJ links two brain networks integrating external (sensory) vs. internal (memory, social-oriented stimuli) information^26^. It is conceivable that a conflict between external instruction (avoid unhealthy food), and internal impulse (approach unhealthy food) increases activity of the rAG in order to solve this conflict. This is consistent with studies showing that the rAG is directly engaged in resolution of stimulus-response conflicts, but also attentional reorientation and response inhibition^32-37^.

Though approach behavior towards unhealthy food pictures decreased, approach behavior towards healthy food pictures remained unchanged. This is in line with previous findings, where decreasing approach behavior towards unhealthy food was the main training effect^4,38^. In our study, decreasing approach tendencies towards unhealthy food was related to decreased brain activation in the rAG, suggesting that training makes avoiding food a less conflicting and more automatic behavior. Further, we found no conclusive evidence for generalization effects. While generalization was repeatedly observed in other contexts (e.g.^8^), results in the obesity context have been mixed^4,38^.

For unhealthy food avoidance, we found increased *post*-training task-related connectivity between the rAG and the dorsal striatum, which is related to stimulus-response learning, executive attention and exerting cognitive control^39-43^. Increased connectivity between the dorsal striatum and rAG was previously related to explicit usage of learned stimulus-response-outcome associations^44^. This could be interpreted as a complementary effect of training. Decreased activity of the rAG was related to more efficient inhibition of automatic reactions after training, possibly through its increased coupling with other brain structures, suggesting that the rAG does not require similar activation strength as *pre* training. This interpretation is supported by resting-state connectivity results, where we found higher *post*-training resting-state connectivity between inhibitory regions in the brain – the bilateral MFG, and left IFG. The left MFG was previously shown to be activated for viewing high caloric food pictures^45^. Changed connectivity between the left and right MFG, structures engaged in response inhibition^28-31^, might therefore be associated with training-induced stronger inhibitory tendencies towards unhealthy food pictures. We also found higher *post*-training connectivity of a reward related region - the nucleus accumbens - and the left IFG, also engaged in response inhibition^46^. This could be associated with increased inhibition of approach response to rewarding stimuli.

In comparison to the alcohol context, where effects of CBM seem to be mediated by altering the rewarding properties of problematic stimuli, underlying neural mechanisms of CBM seem to differ in the food context. This might be explained by differences in the duration of training. Changes in stimuli evaluation and reward-related areas, as found in the alcohol context, are more of a reflective process, thus requiring longer training. However, a previous intervention study in obese individuals, applying food response inhibition training over a four week period, found decreased brain activations in the insula, inferior parietal lobe and putamen^47^. It seems that a longer training in the food context elicits changes in brain areas similar to those in our study. Hence, an alternative explanation of discrepancies between results in the food and alcohol context is that training in the food context works in a different way, possibly because food biases may depend in a different neural network.

It is important to consider limitations of this study. Firstly, sample size was moderate (33 participants). Secondly, CBM effects did not translate to the picture-sorting task, which aimed to assess explicit evaluations of food stimuli. This could point towards CBM training inhibition rather than affecting evaluation processes. This lack of transfer from implicit training to explicit evaluation is in correspondence with the lack of neuronal effects in valuation areas. Further, our training only focused on healthy and unhealthy categories without additional divisions, e.g. into sweet and savory. This may decrease the sensitivity of our analyses and may be related to lack of effects on the picture-sorting task. Also, we compared approach and avoidance tendencies between healthy and unhealthy food images, not including neutral images of non-food objects. Further, due to ethical reasons, we did not train obese participants to approach unhealthy and avoid healthy food cues. Importantly, we do not show effects of training on food intake, which was not assessed in this study. This is a main goal of CBM studies and has only been investigated to a small degree^17,48^. Future studies should implement CBM interventions in real-life settings, assessing its impact on eating behavior.

However, our study provides a basis for future studies on food approach bias modification, which could focus on specifically strengthening inhibitory control, especially regarding unhealthy food. Additionally, showing that already one CBM session can modify approach tendencies in the laboratory context is very promising. In conclusion, we were able to show that obese individuals have automatic approach tendencies towards food. We further present a possibility to retrain and decrease approach tendencies, especially towards unhealthy foods, and give insight into underlying neural mechanisms. This study could constitute a basis for intervention programs utilizing similar behavioral paradigms. We suggest that these studies implement longer training periods, similar to ones used with alcohol-dependent patients. Further, trainings should specifically aim at strengthening inhibition, specifically towards unhealthy food, rather than encouraging approach towards healthy food, as often done in weight loss programs. Additionally, by showing neural correlates of CBM, our results contribute to possible brain stimulation research focusing on decreasing approach bias towards food.

## Supporting information

## Acknowledgments

Authors would like to thank Jöran Lepsien for valuable input concerning fMRI analysis methods, and Laura Rehberger, Mandy Jochemko, Nicole Pampus and Anke Kummer for help with participants’ recruitment and data acquisition.

Author contributions

NM, FM, and AH designed the research. NM and FM performed the research. All authors analysed the data and wrote the paper.

## Notes

**Financial support** NM was supported by MaxNetAging Research School. The work of FM and AH is supported by the Integrated Research and Treatment Center AdiposityDiseases at the University of Leipzig. The work of AH and AV is supported by the Collaborative Research Centre 1052 ‘Obesity Mechanisms’, subproject A5, at the University of Leipzig.

