## Supplementary material for "Unhealthy Yet Avoidable – How Cognitive Bias Modification Alters Behavioral And Brain Responses To Food Cues In Obesity"

**Supplementary materials and methods**

1. Questionnaire measures

In our study we included the three factor eating questionnaire (TFEQ[1]), assessing eating behavior on three dimensions: cognitive restraint, disinhibition and hunger. The behavioral inhibition/activation system (BIS/BAS[2]) was used to evaluate how a person's behavior is driven by reward and punishment. Questionnaires were used as baseline comparison of groups included in the study. Distribution of all questionnaire measures was normal or close to normal, and equal variance between groups was assumed for all the tests.

2. Preprocessing

AAT-fMRI and rsfMRI data were preprocessed in a similar fashion. Data were preprocessed and statistically analyzed using FMRIB Software Library 5.0.8 (FSL, The University of Oxford, Oxford, United Kingdom [3], SPM 12 revision 6225 (Wellcome Department of Cognitive Neurology, London, United Kingdom), Analysis of Functional NeuroImages version 17.0.04n (AFNI, [4]), Advanced Normalization Tools (ANTs [5]) and MATLAB R2012b (The MathWorks, Inc., Natick, Massachusetts, United States). Firstly, to enable further preprocessing steps, high resolution anatomical images were skull-stripped using FSL's brain extraction tool [6] and SPM 12 segmentation tool. Functional data were motion corrected using McFLIRT [7], fieldmap corrected and registered to high resolution anatomical images (FLIRT, boundary based registration [7-9], slice-timing corrected, and smoothed with a 6mm FWHM Gaussian kernel (not rsfMRI data used for connectivity analysis [10] using FSL's FEAT. To ensure that motion- and physiological noise-related artefacts were removed from the functional time-series, we used independent component analysis automatic removal of motion artefacts (ICA AROMA, [11]) toolbox on the time-series. Further, we regressed out the signal in white matter and cerebrospinal fluid from the functional data. Then, anatomical images were normalized to a 3mm MNI template using ANTs. Using transformation information from the previous registration steps, functional images were registered to the 3mm MNI template using ANTs. Prior to statistical analysis on an individual level, AAT-fMRI data were high-pass filtered with a filter of 128s (SPM). Prior to connectivity analyses, resting state data were high pass filtered (FSL; $\sigma$=22).

3. GLM1 details

On a single-subject level we entered pre and post trials into a general linear model. This resulted in 16 different regressors over 2 sessions. 4 regressors for the picture presentation period pre training (healthy_pull, healthy_push, unhealthy_pull, unhealthy_push), 4 regressors for the zooming period pre training (corresponding to four different types of trials) and a similar set of 8 regressors for the post phase. For the picture presentation period, onsets of the regressors were time-locked to the picture presentation, and event duration was equal to the reaction time. This variable epoch model was described as the most appropriate for reaction time tasks [12]. The onsets of the zooming period regressors were time-locked to the end of the picture presentation period, and the durations were set to 750ms.

4. Definition of seeds

To investigate task-unrelated connectivity differences caused by CBM, we defined a number of seeds directly related to reward processing, visual food stimuli processing and inhibitory control. This was done in order to test our hypotheses of reward vs. inhibitory mechanisms involved in the CBM. The following seeds were included in our study: the medial and the left and right dorsolateral prefrontal cortex (mPFC, dlPFC, coordinates from: [13]), the left and right amygdala and nucleus accumbens (Amy, NAcc, coordinates from pickatlas [14] and the left middle frontal gyrus (MFG, coordinates from [15]). The mPFC, amygdala and the nucleus accumbens were previously shown to be engaged in approach-avoidance tendencies and are widely accepted reward-related brain regions [16]. The mPFC is widely accepted as the brain's valuation center [17], the amygdala is important for Pavlovian learning and formation of emotional memories [18, 19], whereas the nucleus accumbens receives and sends dopaminergic projections in response to rewarding stimuli [18-21]. The dlPFC was previously related to approach-avoidance tendencies and is an inhibitory brain region [22, 23]. Lastly, the left MFG is a region preferentially activated for viewing high versus low caloric food stimuli [15].

**Supplementary results**

1. Questionnaire results

There were no baseline group differences for all questionnaire variables (smallest p = .082, Table S2).

Table S1: Sample characteristics of the training and sham-training groups; p-values reflect significance of group differences

| | Training Group | No-Training Group | p-value/t(31) value (unless otherwise indicated) | Effect size \|d\| (unless otherwise indicated) |
|---|---|---|---|---|
| | Mean/SD (unless otherwise indicated) | | | |
| n | 17 | 16 | | |
| Sex | 11 ♀. 6 ♂ | 7 ♀. 9 ♂ | 0.227/$\chi^2$=1.460 | ϕ=0.043 |
| Age [years] | 28/5 | 31/4 | **0.027/2.314** | 0.663 |
| BMI [kg/m$^2$] | 35.57/4.63 | 36.95/7.63 | 0.530/0.635 | 0.219 |
| Hunger [VAS cm; not hungry – hungry] | 2.31/1.81 | 2.73/1.91 | 0.534/0.629 | 0.226 |
| Tiredness [VAS cm; not tired – tired] | 4.31/2.60 | 4.00/2.30 | 0.726/-0.354 | 0.126 |
| Mood [VAS cm; in a bad mood – in a good mood] | 7.69/1.58 | 8.20/1.26 | 0.711/=0.374 | 0.357 |

Table S2: Questionnaire differences between the training and no-training groups; p-values reflect significance of group differences

| | Training Group | No-Training Group | p-value/t(31) value | Effect size \|d\| |
|---|---|---|---|---|
| | Mean/SD | | | |
| TFEQ cognitive control | 8.11/6.12 | 6.63/4.01 | 0.417/-0.822 | 0.286 |
| TFEQ disinhibition | 9.29/3.16 | 7.19/3.56 | 0.082/-1.800 | 0.624 |
| TFEQ hunger | 6.94/4.35 | 5.75/3.44 | 0.391/-0.869 | 0.303 |
| BIS | 20.53/4.61 | 19.94/5.23 | 0.732/-0.345 | 0.120 |
| BAS drive | 12.00/2.06 | 11.62/2.63 | 0.651/-0.457 | 0.161 |
| BAS fun seeking | 13.18/2.16 | 12.38/1.89 | 0.267/-1.131 | 0.394 |
| BAS reward responsivity | 16.76/1.92 | 16.75/1.39 | 0.980/-0.025 | 0.006 |

Table S3. Ratings of healthy and unhealthy images used in the AAT paradigm for healthiness and liking on a scale from 0 to 10

| Image category | Scale | Training Group | Sham-training Group | p-value/t(31)-value | Effect size \|d\| |
|---|---|---|---|---|---|
| | | Mean/SD | | | |
| Unhealthy | Liking | 6.11/0.90 | 5.95/1.34 | 0.692/-0.399 | 0.140 |
| | Healthiness | 1.81/0.78 | 1.76/1.21 | 0.885/-0.146 | 0.049 |
| Healthy | Liking | 7.64/0.71 | 7.44/1.17 | 0.555/-0.596 | 0.207 |
| | Healthiness | 8.67/0.67 | 8.73/0.62 | 0.775/0.288 | 0.093 |

Table S4. Picture-sorting task – ratings of healthy and unhealthy images on healthiness and liking on a scale from 0 to 10. We observed no significant group differences.

| Image Category | Scale | Training Group | | Sham-training Group | | Image category * time * group interaction | Effect size $\eta^2_p$ |
|---|---|---|---|---|---|---|---|
| | | Pre | Post | Pre | Post | | |
| | | Mean/SD | | | | p-value/F(1,31)-value | |
| Liking | Unhealthy | 5.76/ 1.17 | 5.37/ 1.57 | 5.97/ 1.82 | 5.39/ 1.94 | 0.975/0.001 | 0.014 |
| | Healthy | 7.24 /0.99 | 7.39/ 1.07 | 7.23/ 1.63 | 7.35/ 1.63 | | |
| Healthiness | Unhealthy | 1.38/ 0.82 | 1.53/ 0.88 | 1.25/ 1.01 | 1.50/ 1.16 | 0.503/0.453 | 0 |
| | Healthy | 8.42/ 0.76 | 8.43 /0.83 | 8.27/ 0.97 | 8.38/ 0.90 | | |
